# Reference Library for Suspect Non-targeted Screening of Environmental Toxicants Using Ion Mobility Spectrometry-Mass Spectrometry

**DOI:** 10.1101/2025.02.22.639656

**Authors:** Devin Teri, Noor A. Aly, James N. Dodds, Jian Zhang, Paul A. Thiessen, Evan E. Bolton, Kara M. Joseph, Antony J. Williams, Emma L. Schymanski, Ivan Rusyn, Erin S. Baker

**Affiliations:** Department of Veterinary Physiology and Pharmacology, Texas A&M University, College Station, Texas 77843, USA; Department of Chemistry, University of North Carolina, Chapel Hill, North Carolina 27599, USA; National Center for Biotechnology Information, National Library of Medicine, National Institutes of Health, Bethesda, MD 20894, USA; Center for Computational Toxicology and Exposure, Office of Research and Development, U.S. Environmental Protection Agency, Research Triangle Park, North Carolina 27711, USA; Luxembourg Centre for Systems Biomedicine, University of Luxembourg, 6 Avenue du Swing, 4367, Belvaux, Luxembourg

**Keywords:** mass spectrometry, ion mobility spectrometry, non-targeted analysis, exposure assessments, environmental analysis

## Abstract

As our health is affected by the xenobiotic chemicals we are exposed to, it is important to rapidly assess these molecules both in the environment and our bodies. Targeted analytical methods coupling either gas or liquid chromatography with mass spectrometry (GC-MS or LC-MS) are commonly utilized in current exposure assessments. While these methods are accepted as the gold standard for exposure analyses, they often require multiple sample preparation steps and more than 30 minutes per sample. This throughput limitation is a critical gap for exposure assessments and has resulted in an evolving interest in using ion mobility spectrometry and MS (IMS-MS) for non-targeted studies. IMS-MS is a unique technique due to its rapid analytical capabilities (millisecond scanning) and detection of a wide range of chemicals based on unique collision cross section (CCS) and mass-to-charge (*m/z*) values. To increase the availability of IMS-MS information for exposure studies, here we utilized drift tube IMS-MS to evaluate 4,685 xenobiotic chemical standards from the Environmental Protection Agency Toxicity Forecaster (ToxCast) program including pesticides, industrial chemicals, pharmaceuticals, consumer products, and per- and polyfluoroalkyl substances (PFAS). In the analyses, 3,993 [M+H]^+^, [M+Na]^+^, [M-H]^-^ and [M+]^+^ ion types were observed with high confidence and reproducibility (≤1% error intra-laboratory and ≤2% inter-laboratory) from 2,140 unique chemicals. These values were then assembled into an openly available multidimensional database and uploaded to PubChem to enable rapid IMS-MS suspect screening for a wide range of environmental contaminants, faster response time in environmental exposure assessments, and assessments of xenobiotic-disease connections.

## Introduction

Ion mobility spectrometry coupled with mass spectrometry (IMS-MS) is a non-targeted analytical technique that has stimulated great interest in both analytical chemistry and exposure science due to its ability to resolve isomeric species and distinguish halogenated chemicals (Foster *et al*., 2022; Ibrahim *et al*., 2016; Kanu *et al*., 2008; Kirkwood-Donelson *et al*., 2023; Luo *et al*., 2020; May and McLean, 2015; Metz *et al*., 2017). IMS separates ions by their shape and charge state in the gas phase through the use of an electric field, buffer gas and sometimes gas flow, depending on the IMS type utilized (*e*.*g*., drift tube, traveling wave, trapped IMS, etc.) (Hinnenkamp *et al*., 2018; Jurneczko *et al*., 2012). In addition to its millisecond analysis speed, most IMS methods also allow for the calculation of collision cross section (CCS) values for each ion detected (Ewing *et al*., 2016; Gabelica *et al*., 2019). CCS values correspond to the collision area between a charged analyte and its neutral collision partner or the buffer gas molecules in the IMS cell. In various studies, CCS values have proven to be highly reproducible (often less than 2% of each other) between laboratories using the same instrumentation, even in complex samples (Stow *et al*., 2017). Additionally, CCS values for the same molecule evaluated with the different IMS methods are often within 2% of each other (Hinnenkamp, *et al*., 2018). Therefore, CCS values are regarded as highly informative data which provide a way to increase confidence in the identification of individual molecules in non-targeted analytical studies of complex substances and mixtures (Burnum-Johnson *et al*., 2019; Dodds *et al*., 2020). Moreover, in IMS-MS studies, ions separated by IMS can be evaluated for their *m/z* value, and if fragmentation is applied, 3-dimensional characterization is possible (CCS, MS and MS/MS) (Dodds and Baker, 2019; Ibrahim *et al*., 2015).

Historically, IMS-MS has been used to study peptides, proteins, carbohydrates and inorganic molecules (May *et al*., 2017). More recently, studies of environmental and industrial chemicals have provided additional CCS information to enable analyses of complex environmental exposures (Aly *et al*., 2022; Belova *et al*., 2021; Celma *et al*., 2020; Hinnenkamp, *et al*., 2018; Picache *et al*., 2019; Zheng *et al*., 2018). These studies have demonstrated the use of CCS libraries in suspect screening of complex substances (Cordova *et al*., 2023; Luo, *et al*., 2020; Luo *et al*., 2021; Roman-Hubers *et al*., 2022a; Roman-Hubers *et al*., 2022b) and in assessing environmental samples (Celma, *et al*., 2020; Valdiviezo *et al*., 2022). The opportunities for further advancing chemical exposure analyses and determining potential adverse effects on human health are therefore rapidly evolving with analytical chemistry advances such as IMS-MS (Escher *et al*., 2017; Escher *et al*., 2020; Scholz *et al*., 2022; Vermeulen *et al*., 2020). However, improved linkages between exposure measurements and potential hazards of chemicals, such as high-throughput toxicity data (Williams *et al*., 2017), are still lacking due to the absence of analytical methods that can be used for rapid analyses of complex samples and mixtures (Barouki *et al*., 2022; Brack *et al*., 2016; Lai *et al*., 2024). Additionally, refinements to characterization of *in vitro* to *in vivo* extrapolations (IVIVE) of individual chemicals (Wambaugh *et al*., 2018; Wambaugh *et al*., 2019) and mixtures (Valdiviezo *et al*., 2021) would therefore benefit from improved analytical methods able to assess diverse chemicals in the environment and their metabolites in biological systems. Fortunately, government agencies have encouraged collaborative analyses of large-scale chemical libraries and mixtures by providing access to their compound libraries (Ulrich *et al*., 2019), enabling opportunities for developing large-scale reference datasets for suspect screening using rapid non-targeted analytical methods such as IMS-MS.

In this study, we follow up on recent studies that have aimed to establish CCS libraries for xenobiotic chemicals (Belova, *et al*., 2021; Celma, *et al*., 2020; Hinnenkamp, *et al*., 2018; Picache, *et al*., 2019) by utilizing drift tube IMS (DTIMS) and nitrogen buffer gas (denoted as ^DT^CCS_N2_) to evaluate chemicals of concern to human health which have also been tested in hundreds of *in vitro* assays by the ToxCast/Tox21 programs (Richard *et al*., 2021; Williams, *et al*., 2017). For these analyses, we utilized an electrospray ionization (ESI) source in both positive and negative mode and an atmospheric pressure chemical ionization (APCI) source in positive mode for more comprehensive molecular coverage of the diverse chemicals. The database created from this study contains CCS values for 2,140 unique chemicals and will be of immediate interest for the chemical analyses of complex samples and environmental mixtures (Phillips *et al*., 2022).

## Materials and Methods

### Sample Preparation

The 4,685 xenobiotic chemical standards analyzed in this study (**Figure 1** and **Tables S1-S2**) were obtained from Evotec (Branford, CT; order #13169), a contractor of the U.S. Environmental Protection Agency’s (US EPA) ToxCast library of chemicals (Richard *et al*., 2016). Chemical selection from a ToxCast library was performed by the U.S. EPA (Dr. Ann M. Richard) based on chemical availability and the purity of the individual compounds at the time of assembly and all standards were supplied at 0.4 mM in pure dimethyl sulfoxide (DMSO). The process for selecting and registering chemicals into the ToxCast library including the description of chemical procurement from commercial suppliers, cataloguing the chemicals, storage and chemical quality control is detailed at (Richard *et al*., 2024; Richard *et al*., 2014). After selection, all chemicals were placed into sealed 384-well plates, shipped, and then stored frozen (−80 LC). The certificate of analysis obtained with the chemical shipment stated all compounds were >90% purity as determined by the supplier. Before experiments, the stock plates were thawed at room temperature, 5 μL of each chemical was then transferred to individual 9 mm amber screw top vials with fused inserts (Ibis Scientific, #4400-FIV2W), and each standard was diluted to 10 μM with 195 μL of 50:50 methanol/water. The high purity (≥99.9%) water, methanol, and acetonitrile used in this study were obtained from Sigma-Aldrich (St. Louis, MO).

**Figure 1.**
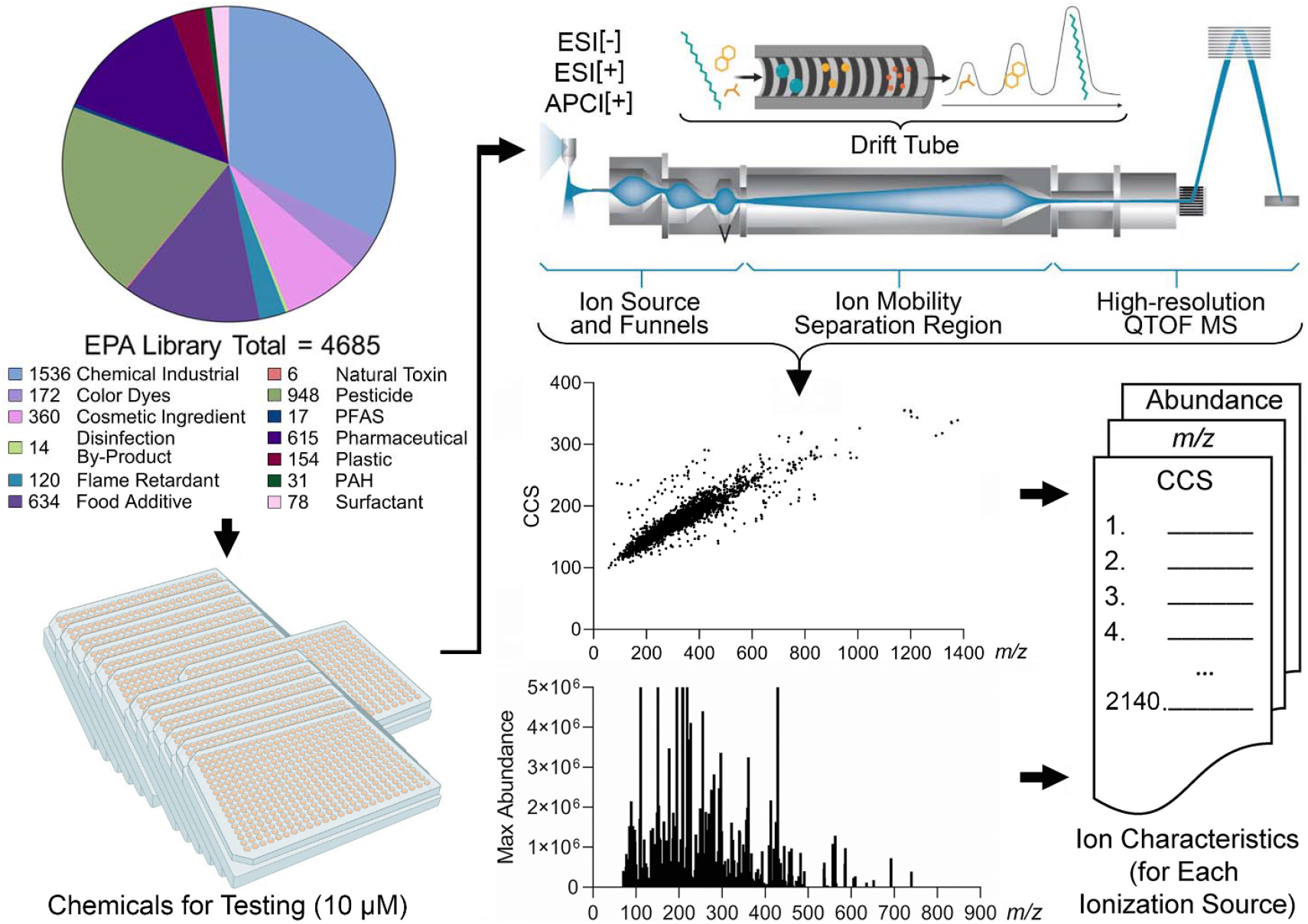
Project Workflow. In this study, we first classified the 4,685 chemical standards into 13 classes. The 4,685 standards were then diluted to 10 μM and analyzed with IMS-MS using ESI[+], ESI[-] and APPI[+]. Feature identification and collision cross section calculations were performed using Agilent IM-MS Browser and Agilent Mass Profiler. CCS values for 2,140 unique chemicals were obtained in the study and 3,993 ions from [M+H]^+^, [M+Na]^+^, [M-H]^-^ and [M+]^+^ ion types.

### Instrumentation and Analysis

DTIMS-MS analyses were performed using the 6560 IM-QTOF MS system (Agilent Technologies, Santa Clara) and the drift tube was filled with nitrogen gas. A liquid chromatography stack (Agilent Technologies: G7116B Column Compartment, G7167B Multisampler, and G7120A High Speed Pump) was used as an autosampler to inject samples into each ionization source at 0.400 mL/min with a constant solvent composition of 50% water and 50% acetonitrile. All chemical standards were ionized using the APCI source in positive ionization mode and the ESI source in both positive and negative ionization mode.

Prior to each experiment, the instrument was tuned and mass calibrated using the Agilent Tune Mix (G2421A/G2432A, Agilent Technologies, Santa Clara, CA). For the IMS analyses, the ions were passed through the inlet glass capillary, focused by a high-pressure ion funnel, and accumulated in an ion funnel trap (Baker *et al*., 2007). Ions were then pulsed into the 78.24 cm long drift tube filled with ∼3.95 torr of nitrogen gas, where they travelled under the influence of a weak electric field (17 V/cm). Ions exiting the drift tube were refocused by a rear ion funnel prior to quadrupole time-of-flight (QTOF) MS analysis and their drift time was recorded. The detailed instrumental settings are listed in **Tables S3-S5**. For each detected ion, ^DT^CCS_N2_ values were calculated using a single-field method which has been detailed previously (Stow, *et al*., 2017). Methanol washes were performed after 24 sample injections to reduce the occurrence of carryover. Additionally, isomeric chemicals were also separated by several standards with greater mass differences to ensure no interferences occurred. Samples from the same plate were run in the same worklist over six days, alternating the ion mode each day. Each chemical was injected six times total, resulting in duplicate results for each ion mode. The average ^DT^CCS_N2_ values having less than 1% difference between replicates are found in **Table S6**.

### Data Analysis and Curation

Agilent IM-MS Browser software (Agilent Technologies) was utilized for all single-field ^DT^CCS_N2_ calculations, initially starting with the Tune Mix ions for use as calibrants. While software programs can assist in calculating the CCS values, every instrumental run was manually examined for the presence of a peak and the software-assigned CCS value was verified. Quality control and confidence in CCS values was determined based on the ^DT^CCS_N2_ duplicate values (GraphPad, 2024) having a percent difference of ≤1 for at least two replicates (Zhang *et al*., 2016). The ^DT^CCS_N2_ and *m/z* values were then plotted to identify potential trendlines. Only ^DT^CCS_N2_ values that fell within 15% difference of the main trendline were determined to be confident identities and listed in the final dataset. This filtering removes small molecules which multimerize in solution, drift through the ion mobility cell as multimers, and then break into monomers following drift analysis, thereby illustrating extremely large CCS values per *m/z* (Baker *et al*., 2023).

Of the 4,685 standards evaluated herein, three chemicals were duplicated in the plates, resulting in 4,682 unique chemicals. The chemicals were assigned into classes using expert judgement of the authors using PubChem “Use and manufacturing” classifications and other pertinent information in the US EPA’s CompTox Chemicals Dashboard. Following ESI-IMS-MS and APCI-IMS-MS analyses, reproducible CCS values resulted for 2,140 unique chemicals. Moreover, due to the various ion types observed ([M+H]^+^, [M+Na]^+^, [M-H]^-^ and [M+]^+·^), the IMS-MS database contains 3,993 CCS values and the corresponding *m/z* ratios.

## Results and Discussion

Many experimentally measured CCS datasets have been published for use in the fields of proteomics, metabolomics, steroids, and xenobiotics (Aly, *et al*., 2022; Belova, *et al*., 2021; Celma, *et al*., 2020; Hinnenkamp, *et al*., 2018; Picache, *et al*., 2019; Zheng, *et al*., 2018), but the number of CCS values is still limited, especially for xenobiotic studies. The purpose of developing a database of CCS values from the EPA’s ToxCast library is therefore to assist in identifying toxicants and improve the assessment of human and environmental risks and exposures. To create the IMS-MS ToxCast database, 4,685 chemical standards were analyzed using the workflow depicted in **Figure 1**. In this workflow, the chemicals were first categorized into classes, diluted, evaluated with IMS-MS, assessed for reproducible CCS values, and then added to a CCS database if they passed all quality control steps stated in the Methods section. For the categorization step, the chemicals were placed into thirteen broad classes based on their structures and use by employing information from the EPA’s ToxCast library (Williams, *et al*., 2017). Because most chemicals had several classifications listed, we used the source/use information in PubChem to determine the final assignments. Overall, the following assignments were made: Natural Toxin (n=6), Disinfection By-Product (n=14), PFAS (n=17), Polycyclic Aromatic Hydrocarbon (PAH, n=31), Surfactant (n=78), Flame Retardant (n=120), Plastic (n=154), Color Dye (n=172), Cosmetic Ingredient (n=360), Pharmaceutical (n=615), Food Additive (n=634), Pesticide (n=948), and Chemical Industrial (n=1,536), where the three duplicates were from the Pharmaceutical, Food Additive, and Chemical Industrial classes. It should be noted that while these classifications are subject to interpretation, the assignment of substances into these classes is commonly used in regulatory science.

For the IMS-MS analyses, each chemical standard was diluted to 10 μM and injected twice into the instrument using ESI[+], ESI[-], and APCI[+] sources. While various ions were formed in the different sources, we specifically evaluated [M-H]^-^ in the ESI[-] analyses, [M+H]^+^ and [M+Na]^+^ in the ESI[+] analyses, and [M+H]^+^ and [M+]^+·^ in the APCI[+] studies. Furthermore, an ion was considered present if the mass error was less than 10 ppm from the expected chemical formula, in addition to the presence of the correct ^13^C isotopic distribution. **Figure 2** shows three example chemicals following ionization using all three ion modes. These chemicals (bosentan, thiabendazole, and triamterene) represent two different chemical classes studied (two pharmaceuticals and a pesticide), each having ions observed in all analysis modes.

**Figure 2.**
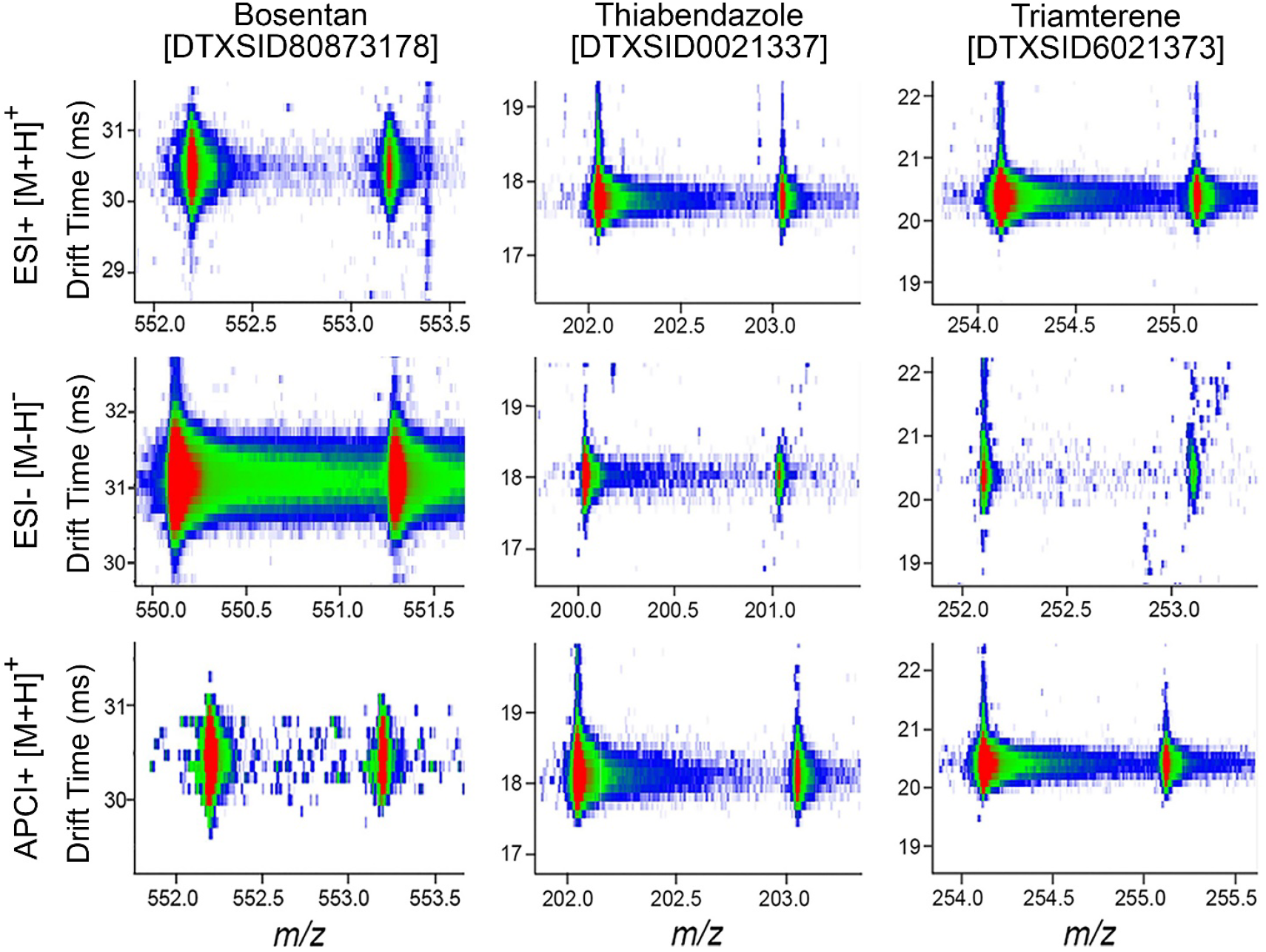
Representative Nested IMS–MS Spectra for Three Test Chemicals. Images illustrate the abundance of the isotopic distribution of the precursor [M] and first ^13^C isotope [M+1] peak for each chemical across the three ionization modes (ESI[-], ESI[+] and APCI[+]). For each chemical shown (Bosentan, Thiabendazole, and Triamterene), *m/z* values are plotted on the x-axis, and drift times on the y-axis. Colors represent abundance from red (highest) to green (intermediate), to blue (lowest). The DTXSIDs are the substance identifier associated with the US EPA’s CompTox Chemicals Dashboard.

For the 4,682 unique chemicals analyzed, ∼45.7% (n=2,140) could be detected in at least one analysis mode (**Figure 3**). This number was not surprising as some chemicals may have degraded or were not amenable to the ionization sources used (such as some of the metals in the ToxCast library (**Table S1**) would need to be run with inductively coupled plasma (ICP)-MS). As expected, differences among chemical classes were observed with respect to what ionization mode was preferable for their detection. For example, of the ion species evaluated, PAHs were primarily detected with APCI[+], while PFAS were identified with ESI[-], due to the structural components of both molecular classes (**Figures 3A** and **3B**). **Figure 3A** breaks down the number of chemicals in each classification detected by each ion source. **Figure 3B** shows the total percentage of unique chemicals detected in every classification and their ion mode for detection. Specifically, no class of chemicals was completely detected in the study, and detectability ranged from less than 20% for “Food Additives” to 75% for both “Color Dyes” and “Pharmaceuticals”. Additionally, ESI[+] provided the majority of identifications (n=1,324), and of this total 426 chemicals were detectable only with ESI[+] (**Figure 3C**). ESI[-] had the second most with n=1,146, and here 634 were detected only with ESI[-]. Finally, APCI[+] had the least amount of chemicals detected at n=771, and only 52 were uniquely detected with this source and polarity. To understand the positive mode detections, we also analyzed which ions were detected using ESI[+] and APCI[+] (**Figure 3D**). For ESI[+], the [M+H]^+^ and [M+Na]^+^ were found to have a similar rate of detection with ∼36% of chemicals in ESI[+] producing both ion types. On the other hand, APCI[+] ion types were abundantly represented by [M+H]^+^, while [M+]^+·^ detection only included 13.1% of the chemicals.

**Figure 3.**
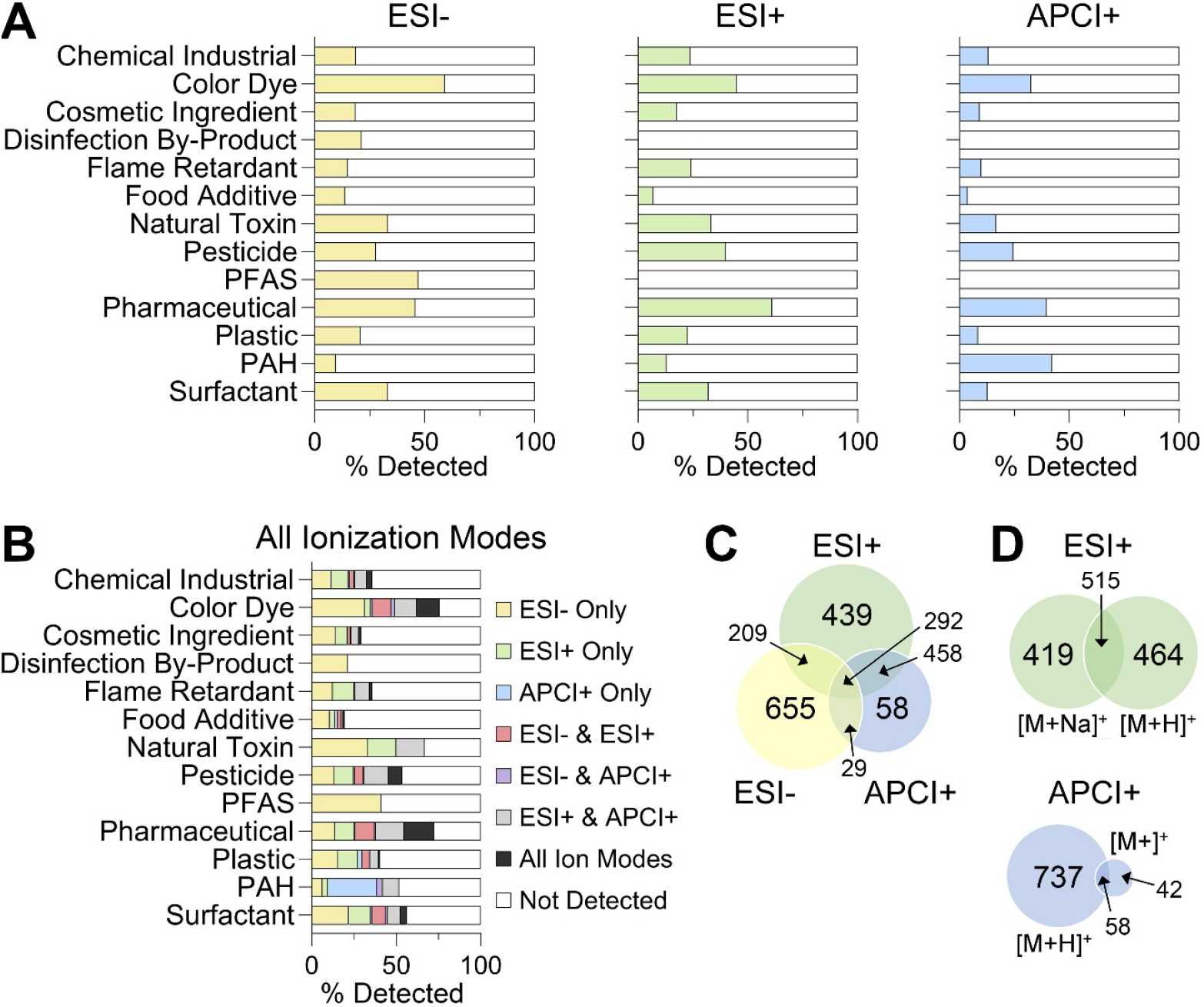
Detection Frequency for Tested Chemicals Across Ionization Modes and Chemical Classes. (A) Percentages of chemicals detected in each class (filled bars) using ESI[-], ESI[+], or APCI[+]. (B) Percentage of chemicals in each class detected by one or multiple ionization modes. (C) Venn diagram illustrating overlap of chemicals detected across the different ionization modes. (D) Venn diagrams showing number of ions detected in ESI[+] and APCI[+].

Because the library of CCS values for chemicals analyzed herein may be used for suspect screening of complex substances and mixtures by other laboratories, we also assessed reproducibility of CCS values. For this examination, we evaluated reproducibility both between the technical replicates in our own analyses (intra-laboratory) and the differences in our values and those published by others (inter-laboratory) using DTIMS and traveling wave IMS (TWIMS) (**Figure 4**). For most chemical classes, over 90% of the substances were within 1% difference. The percent difference in CCS values among the duplicate injections of each chemical were evaluated by chemical class (**Figure 4A**) and ionization mode **(Figure 4B**). Box and whisker plots show the distribution of the deviation among technical replicates, and the pie charts show the fraction of the substances with values under the generally accepted cutoff of 1%. Interestingly, “Natural Toxins” and “PFAS” had the lowest number of retained chemicals using these cutoffs with 14% and 30% removed, however, these were among the classes with the fewest number of chemicals represented in the library. With respect to ionization mode analysis, the overwhelming majority (97%) of the chemicals detected with either ESI[-] or ESI[+] had excellent reproducibility with differences in CCS ≤1%. Even though the reproducibility of CCS values was less robust with APCI[+], 91% were below the threshold of 1%. Of the 3,948 CCS values for the different ion types evaluated, only 171 were excluded from the final database because they exceeded 1% difference, resulting in 3,993 ions in the final database (**Table S6**).

**Figure 4.**
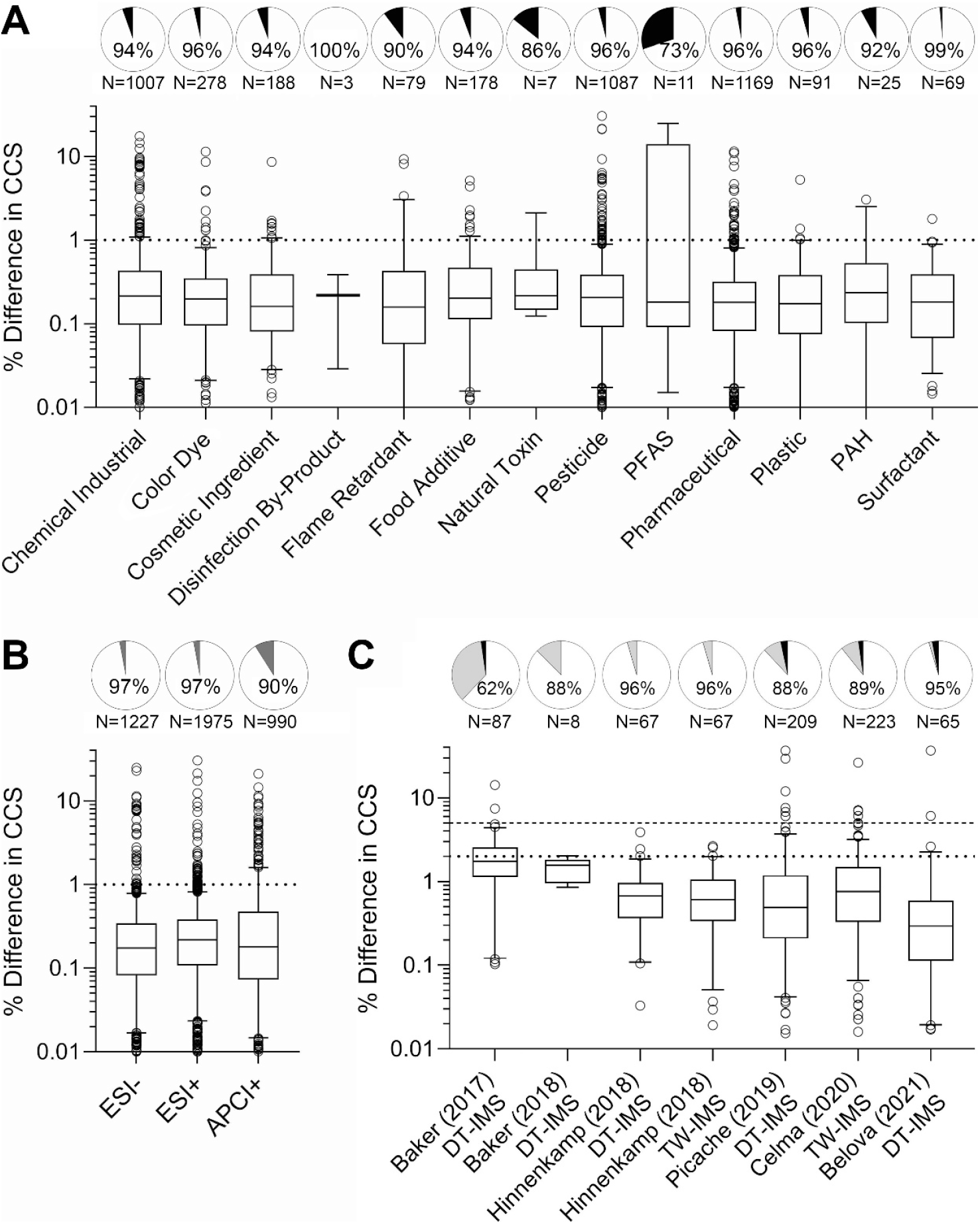
Reproducibility of Collision Cross Section (CCS) Values. CCS reproducibility for the technical replicates examined in this study was assessed by (A) chemical class and (B) ionization mode (ESI[-], ESI[+], and APCI[+]). (C) CCS reproducibility between this study and other published large datasets that contained overlapping chemicals. In panels A and B, the pie charts show the fraction of the chemicals falling below a 1% threshold (white) and above 1% threshold (black), which were filtered out. The total number of chemicals is also noted below each pie chart. In panel C, the pie charts show the fractions below 2% (white), between 2 and 5% (gray), and above 5% (black). The box and whiskers plots in each panel show the distribution of the values for each detected chemical (boxes are interquartile range, the line is median, the whiskers are 5-96 percentile, and the points are outliers). Horizontal lines indicate thresholds used in each analysis.

Interlaboratory reproducibility was also determined in the study using data from six other suspect screening CCS libraries (Belova, *et al*., 2021; Celma, *et al*., 2020; Hinnenkamp, *et al*., 2018; May, *et al*., 2017; Picache, *et al*., 2019; Zheng *et al*., 2017; Zheng, *et al*., 2018) (**Figure 4C and Tables S7-S11**). Our database containing 3,993 CCS values represents a great expansion on important toxicants as most of the other library values are endogenous metabolites. Furthermore, in overall CCS reproducibility assessments across laboratories (*i*.*e*., % difference), it is often considered acceptable if the difference in reported values is ≤2% to account for instrument variability. A 5% difference was also used for this comparison to consider any experimental differences. Of the chemicals that could be cross compared (from 8 to 221 depending on publication), over 87% of the CCS values had ≤2% difference and only ∼2% of the compared chemicals fell above the 5% threshold. These small errors show that CCS is an intrinsic chemical property, and only slight errors occur due to differences in electric field application, CCS calibration, and pressure or temperature fluctuations in the instrumentation. Overall, these comparisons provide us with confidence in our database as all values included fell below the 1% difference threshold and we had very few values outside of the 5% difference when compared to other databases.

To finalize this study, we attempted to determine reasons for lack of detection of some of the tested compounds with IMS-MS and the ESI and APPI sources. The main two factors considered were stability of the chemicals in solution (Richard, *et al*., 2016), and amenability of a chemical for detection using a particular ionization source and mode (*e*.*g*., ESI[-] and ESI[+]) (Charest *et al*., 2024; Lowe *et al*., 2021). For the chemical stability analysis, we reasoned that chemicals that lack long-term stability in solution will not be detected due to potential degradation prior to analysis. To make this comparison, we were able to obtain information on long-term stability of the ToxCast library from the U.S. EPA and categorize them using this information as “stable” (*i*.*e*., no appreciable degradation) or “unstable” (chemical not detected in the analysis of a stock solution after 4 months of being stored at room temperature). **Figure 5A** divides the chemicals into 4 categories based on the stability and whether they were detected in at least one ionization mode. The overall number of chemicals included in this analysis was less than the total based on the availability of stability data – out of the 4,682 unique chemicals tested, data was only available for 3,659. Of these, 37.6% were both detected and stable, while 26.6% were neither detected nor stable. This result was highly significant (p<0.0001 using Fisher’s exact test) indicating that stability of the chemicals was a considerable contributor to our ability to determine their CCS value. For the chemical amenability analysis, we reasoned that chemicals that could not be ionized will not be detected. Using previously published predictive chemical amenability analyses with LC-MS (Lowe, *et al*., 2021), we determined which chemicals were “amenable” to ESI[-] or ESI[+] ionization (**Figures 5B** and **5C)**. For each analysis, 4,591 and 4,542 compounds out of the total of 4,682 analyzed could be predicted for each ionization mode. In the ESI[-] analysis, 71.8% of the chemicals were predicted as non-amenable. This is an indication that GC-IMS-MS should be a future goal for analysis of these chemicals. Additionally, of the amenable chemicals, 59.2% were detected in the ESI[-] evaluations. In the ESI[+] analysis, 57% of the chemicals were predicted as non-amenable and for the amenable chemicals, 55.8% were detected. These analyses demonstrated that amenability was also a major contributing factor to our ability to derive CCS values (p<0.0001 using Fisher’s exact test for both ionization modes) as many chemicals were predicted not amenable. Of additional note, all chemicals were tested at 10 μM and there was no concentration response performed in this study. Thus, it could not be determined if the concentration of the standard was below the limit of detection or if the chemical had degraded over time to a lower concentration. As mentioned previously, each data file was analyzed manually for its specified chemical. This was done to ensure the most accurate analysis and incorporate expert opinions in cases of discrepancies regarding features like saturation, isomers, low abundances, and n-mer streaking (*e*.*g*., where unstable multimers break into monomers during instrumental analysis). However, this does introduce bias into the experiment.

**Figure 5.**
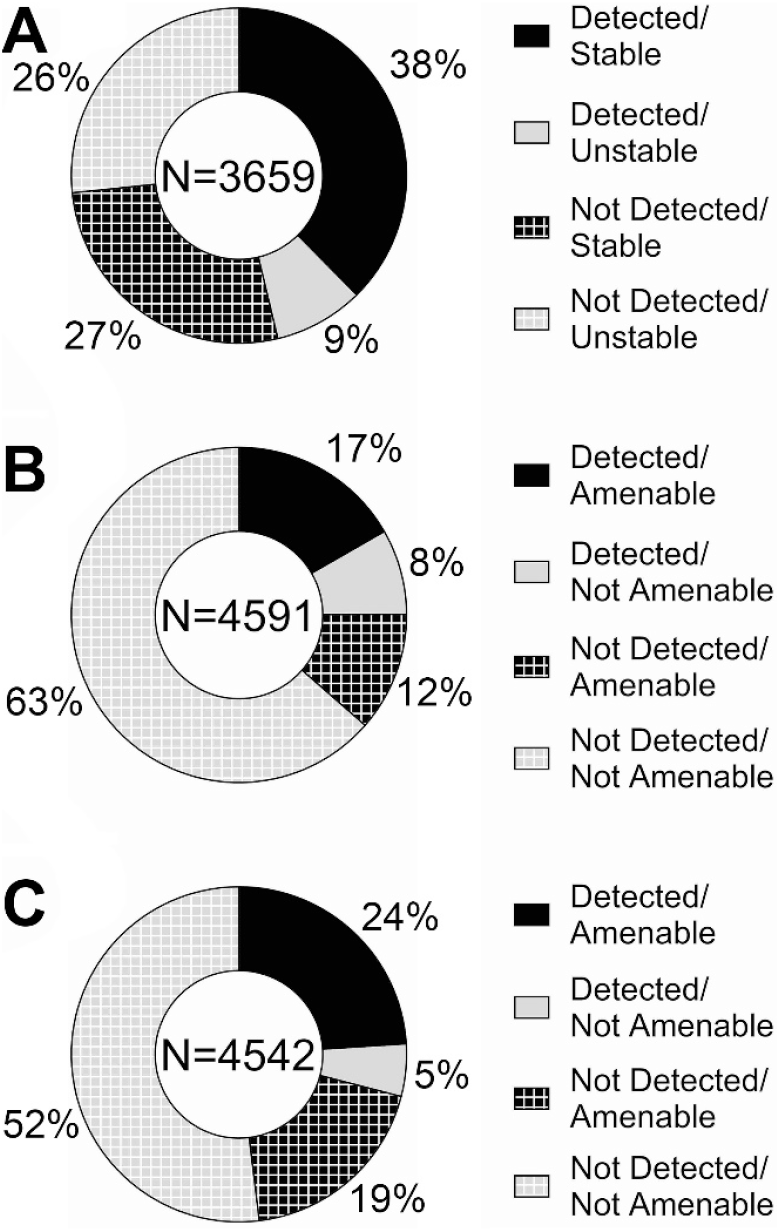
Analysis of Chemical Stability and Amenability as Factors Influencing Detection. Donut plots show fractions of chemicals based on whether they were detected and their (A) stability, (B) amenability to detection with ESI[-], and (C) amenability for detection with ESI[+]. The total number of chemicals included in each analysis is shown at the center of each plot.

## Conclusions

In this study, we utilized IMS-MS to develop a multidimensional database containing experimentally measured CCS and *m/z* values for 2,140 unique chemicals from the U.S. Environmental Protection Agency ToxCast program. The 4,685 chemicals evaluated covered thirteen classes including Natural Toxin (n=6), Disinfection By-Product (n=14), PFAS (n=17), Polycyclic Aromatic Hydrocarbon (PAH, n=31), Surfactant (n=78), Flame Retardant (n=120), Plastic (n=154), Color Dye (n=172), Cosmetic Ingredient (n=360), Pharmaceutical (n=615), Food Additive (n=634), Pesticide (n=948), and Chemical Industrial (n=1,536). To create the database, each chemical standard was assessed twice with ESI[+], ESI[-] and APCI[+] followed by IMS-MS analyses. Chemicals were then identified based on expected *m/z* values for [M+]^+·^, [M+H]^+^, [M+Na]^+^ and [M-H]^-^ ion types. From these measurements, only chemicals having CCS values ≤1% error in the duplicate assessments were retained as high confidence identifications. This resulted in 2,140 unique chemical entries and 3,993 ion types. It was also noted that ESI[-] and ESI[+] worked for the largest majority of the chemicals, however APCI[+] covered a unique subset and provided value for certain chemical classes including the PAHs. Furthermore, comparison of the CCS database values with 6 other laboratories showed ∼88% of the overlapping values were within 2% error, indicating interlaboratory reproducibility. Thus, new toxicant knowledge in this chemical library expands on our analytical capabilities and provides a tool for the scientific community to enable rapid IMS-MS screening of a wide range of xenobiotic chemicals, ultimately decreasing response time of exposure assessments and the ability to probe novel disease linkages.

## Author Contributions

The manuscript was written through contributions of all authors, and all authors have given approval to the final version of the manuscript.

## Acknowledgements

This work was funded by a grant from the National Institute of Environmental Health Sciences (P42 ES027704) and a cooperative agreement with the Environmental Protection Agency (STAR RD 84003201). ELS acknowledges funding support from the Luxembourg National Research Fund (FNR) for project A18/BM/12341006. JZ, PAT, and EEB were supported by the National Center for Biotechnology Information of the National Library of Medicine (NLM), National Institutes of Health.

